# Title: Multi-Scale LM/EM Neuronal Imaging from Brain to Synapse with a Tissue Clearing Method, Sca*l*eSF

**DOI:** 10.1101/2021.04.02.438164

**Authors:** Takahiro Furuta, Kenta Yamauchi, Shinichiro Okamoto, Megumu Takahashi, Soichiro Kakuta, Yoko Ishida, Aya Takenaka, Atsushi Yoshida, Yasuo Uchiyama, Masato Koike, Kaoru Isa, Tadashi Isa, Hiroyuki Hioki

## Abstract

The mammalian brain is organized over sizes that span several orders of magnitude, from synapses to the entire brain. Thus, a technique to visualize neural circuits across multiple spatial scales (multi-scale neuronal imaging) is vital for deciphering brain-wide connectivity. Here, we developed this technique by coupling successive light microscope/electron microscope (LM/EM) imaging with an ultrastructurally-preserved tissue clearing method, Sca*l*eSF. Our multi-scale neuronal imaging incorporates 1) brain-wide macroscopic observation, 2) mesoscopic circuit mapping, 3) microscopic subcellular imaging, and 4) EM imaging of nanoscopic structures, allowing seamless integration of structural information from the brain to synapses. We applied the technique to three neural circuits of two different species, mouse striatofugal, mouse callosal, and marmoset corticostriatal projection systems, and succeeded in the simultaneous interrogation of their circuit structure and synaptic connectivity in a targeted way. Our multi-scale neuronal imaging will significantly advance the understanding of brain-wide connectivity by expanding the scales of objects.

## MAIN TEXT

### Introduction

Connectomics, a description of a wiring diagram of the nervous system, is fundamental for understanding of how the neural circuits process information and generate behavior (*1, 2*). The mammalian brain contains a heterogeneous mixture of billions of neurons with trillions of synapses. Neurons elaborate highly specialized processes that can be over a meter in length for transmitting and receiving information, whereas synapses that connect neurons to one another are several hundred nanometers in size. Hence, the imaging scale required for deciphering brain-wide connectivity of mammalian brains is more than several orders of magnitude (*3*).

Electron microscopy (EM) provides an unparalleled resolution to trace nanometer-thin neuronal processes and identify a synapse unambiguously. Recent advances in volume EM, such as serial block-face scanning EM (SBF-SEM), focused ion beam milling and SEM (FIB-SEM), automated tape-collecting ultramicrotomy (ATUM) with SEM (ATUM-SEM), transmission-mode SEM (tSEM), and transmission EM (TEM) camera array (TEMCA), have enabled us to see ultrastructure within a significant volume of brain, opening up the possibility of assembling a connectome of a mammalian brain (*4, 5*). However, current analysis has been limited to small volumes of tens to hundreds of micrometers in extent (*6–9*).

Fluorescence light microscopy (LM) coupled with genetic labeling methods allows tracking of neuronal processes over long distances to assemble mesoscale connectomic maps for the mouse cerebral cortex and thalamus (*10, 11*) and reconstruct individual neurons with subcellular resolution (*12, 13*). Of particular note, tissue clearing techniques have drastically improved the depth-independent observation of biological samples with fluorescence LM, facilitating connectomic analysis with the scales from the macroscopic/brain to microscopic/subcellular level (*14–16*). However, despite its fundamental advances in spatial resolution (*17*), the resolution of LM does not match the size of a synapse that defines neuronal connectivity. Indeed, axodendritic contacts identified by LM observation are only partially predictive of whether synapses are actually formed (*18, 19*). Importantly, a synapse, which consists of presynaptic membrane, postsynaptic membrane, and a synaptic cleft (chemical synapses) or a neuronal gap junction (electrical synapses), is defined by EM observation (*20, 21*).

Here, we developed a technique to decipher brain-wide connectivity across multiple spatial scales by coupling successive LM and EM (LM/EM) imaging with a tissue clearing technique (multi-scale LM/EM neuronal imaging). To achieve the imaging, we developed a glutaraldehyde (GA)-resistant tissue clearing technique, Sca*l*eSF. We further implemented LM/EM dual labeling that remained stable in the clearing protocol. We applied this technique to mouse striatofugal and marmoset corticostriatal projection systems, and succeeded in the simultaneous interrogation of their circuit structure and synaptic connectivity. In addition, we took advantage of the fact that our developed imaging system permitted high-speed LM imaging of substantial tissue volume at high-resolution followed by subsequent EM observation to capture scarce synaptic contacts with nanoscale resolution formed by brain-wide connectivity. We identified and tracked mouse callosal inputs onto parvalbumin (PV)-positive neocortical interneurons in a targeted way across multiple spatial scales. Our multi-scale neuronal imaging will significantly advance the deciphering of brain-wide connectivity and extend the current comprehensive connectomic analysis.

### Results

#### ScaleSF is a tissue clearing method for multi-scale LM/EM neuronal imaging

Multi-scale LM/EM neuronal imaging requires a technique for tissue clearing that achieves a high level of preservation of ultrastructure and fluorescence signals while simultaneously maintaining potent clearing capability (clearing-preservation spectrum). Of proliferating tissue clearing techniques, an aqueous tissue clearing method, Sca*l*eS, occupies a distinctive position with its effective clearing-preservation spectrum (*22*). However, the clearing protocol of Sca*l*eS, sequential 12 hr incubation in six solutions at 37°C, might lead to less-than-optimal preservation of ultrastructure. Although Sca*l*eSQ(0) is formulated for rapid clearance of brain slices without lipid-extracting detergents, a considerable expansion in sample volume is observed after the treatment (*22*), potentially resulting in morphological artifacts. With the goal of minimizing processing time and changes in sample volume, we developed Sca*l*eSF as an isometric and rapid clearing protocol by modifying the clearing procedure of Sca*l*eS (Fig. 1).

**Fig. 1.**
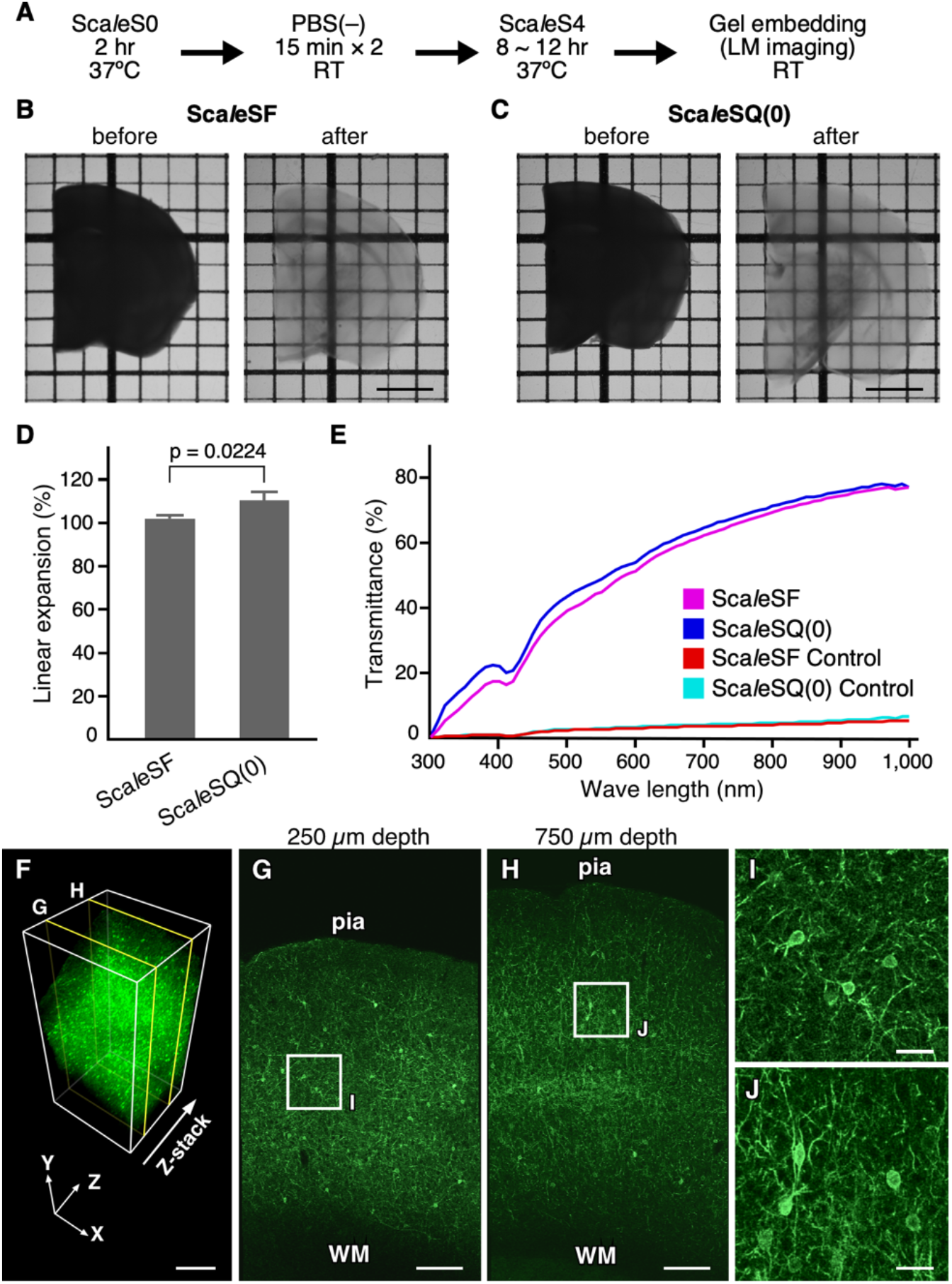
Sca*l*eSF is an isometric and rapid optical clearing method. (**A**) The schedule for tissue clearing with Sca*l*eSF. (**B**, **C**) Transmission images of 1-mm-thick brain slices before (left) and after (right) treatment with Sca*l*eSF (B) and Sca*l*eSQ(0) (C). The grid interval is 1 mm. (**D**) Change in size of brain slices after Sca*l*eSF and Sca*l*eSQ(0) treatment (n = 3, Sca*l*eSF; n = 4, Sca*l*eSQ(0); *t* = 3.261, *df* = 5, *P =* 0.0224, two-tailed unpaired Student’s t-test). Error bars represent SDs. (**E**) Transmission curves of the control, Sca*l*eSF-, and Sca*l*eSQ(0)-treated mouse brain slices (n = 3 brain hemispheres each). (**F**) Three-dimensional volume rendering of the cerebral cortex of a PV-FGL mouse cleared with Sca*l*eSF. In the PV-FGL mouse, somatodendritic membrane-targeted EGFP expression is driven by a parvalbumin promoter. (**G**, **H**) *xy* images in (F) at the depths of 250 μm (G) and 750 μm (H). (**I**, **J**) Enlarged views of rectangles in (G) and (H). pia, pia mater; WM, white matter. Scale bars: 2 mm in (B, C), 500 μm in (F), 200 μm in (G, H), and 40 μm in (I, J).

The clearing protocol of Sca*l*eSF requires the sequential incubation of brain slices in three solutions, Sca*l*eS0 solution, phosphate buffer saline (PBS), and Sca*l*eS4 solution, for a total of 10.5– 14.5 hr (Fig. 1A). Cleared brain slices were embedded in agarose gel dissolved in Sca*l*eS4D25(0) solution (Sca*l*eS4 gel) (*23*). Sca*l*eSF treatment rendered 1-mm-thick mouse brain slices transparent with a similar degree of transparency as that yielded with Sca*l*eSQ(0) (Fig. 1B and C). While a modest expansion in sample sizes was observed after Sca*l*eSQ(0) treatment (linear expansion: 110.7 ± 4.1%) (Fig. 1C and D), the final sizes of brain slices cleared with Sca*l*eSF were approximately 100% of that of the original (linear expansion: 102.5 ± 1.3%) (Fig. 1B and D) after transient shrinkage and expansion (Fig. S2A). The transmission curves of 1-mm-thick mouse brain slices showed that Sca*l*eSF cleared brain slices in a manner comparable to Sca*l*eSQ(0) (Fig. 1E). Although tissues cleared with the original Sca*l*eS protocol can be stably stored in Sca*l*eS4 solution (*22*), brain slices cleared with Sca*l*eSF gradually expanded during storage in the solution (Fig. S2B). This expansion could be controlled by embedding the slices in Sca*l*eS4 gel while still maintaining transparency of the cleared slices (Fig. S2B and C). Thus, Sca*l*eSF is an isometric tissue clearing method with comparable clearing capability to that of Sca*l*eSQ(0).

The fluorescence preservation and clearing capability of Sca*l*eSF were assessed with the brain slices of transgenic mice expressing somatodendritic membrane-targeted enhanced green fluorescent protein (EGFP) in PV-positive neurons (PV-FGL mice) (*24*). The three-dimensional image acquisition of slices collected from the cerebral cortex of the mice was performed with a confocal laser scanning microscope (CLSM) (Fig. 1F–J). The cleared brain slices were placed in a customizable 3D-printed chamber (Fig. S1). The high resolution of the three-dimensional image was demonstrated by *xy* images obtained at different depths (Fig. 1G–J): EGFP targeting of the somatodendritic plasma membrane was discernable even at the depths of 250 μm and 750 μm (Fig. 1I and J), indicating the preservation of both fluorescence signals and membrane structures as well as potent clearing capability of Sca*l*eSF technique.

Fixatives containing glutaraldehyde (GA) improve the preservation of ultrastructural morphology (*25*). However, it remains unclear how GA affects tissue clearing performance and ultrastructural preservation in optically cleared tissues. First, we tested the effects of GA on the clearing capability and isometricity of Sca*l*eSF. Remarkably, Sca*l*eSF treatment rendered GA-fixed brain slices transparent without the shrinkage or expansion of their final sizes, albeit less efficiently transparent in brain slices fixed with high concentrations of GA (1% and 2%) (Fig. 2A–C). Then, we examined the effects of GA on ultrastructural preservation in brain slices cleared with Sca*l*eSF (Fig. 3). To this end, Sca*l*eSF-treated mouse brain slices that had been fixed with GA were restored by washing with PBS (deSca*l*ing) (*26*), and synaptic ultrastructure in the cerebral cortex was imaged by TEM. GA improved ultrastructural preservation even in the brain slices cleared with Sca*l*eSF (Fig. 3A). Raising the concentration of GA in fixatives increased the membrane integrity of presynaptic and postsynaptic structures in the cleared slices (Fig. 3A_1_–A_5_). Scoring the ultrastructural preservation by the membrane continuity of presynaptic terminals demonstrated that, at its low concentration (0.02%), GA improved ultrastructural preservation in the cleared slices to an extent comparable to that in the control slices fixed with paraformaldehyde (PFA) (Fig. 3B). We also noticed that the clearing protocol of Sca*l*eSF failed to fully preserve synaptic ultrastructure (Fig. 3, A_6_, A_7_, and B). The GA-mediated ultrastructural preservation was more dramatic in the brain slices obtained from marmosets (Fig. 3C). Without GA, the membrane integrity of the presynaptic and postsynaptic structures was severely degraded after clearing with Sca*l*eSF (Fig. 3C_1_ and C_4_). By contrast, Sca*l*eSF-treated brain slices fixed with 4% PFA containing 0.2% or 1% GA showed nearly complete contiguous membrane integrity (Fig. 3C_2_–C_6_). We also found that an alternative epoxy resin and a different embedding method were compatible with Sca*l*eSF-treated brain slices (Fig. S3). Collectively, these results indicate that Sca*l*eSF is an isometric, rapid, and GA-resistant tissue clearing method that permits multi-scale LM/EM neuronal imaging.

**Fig. 2.**
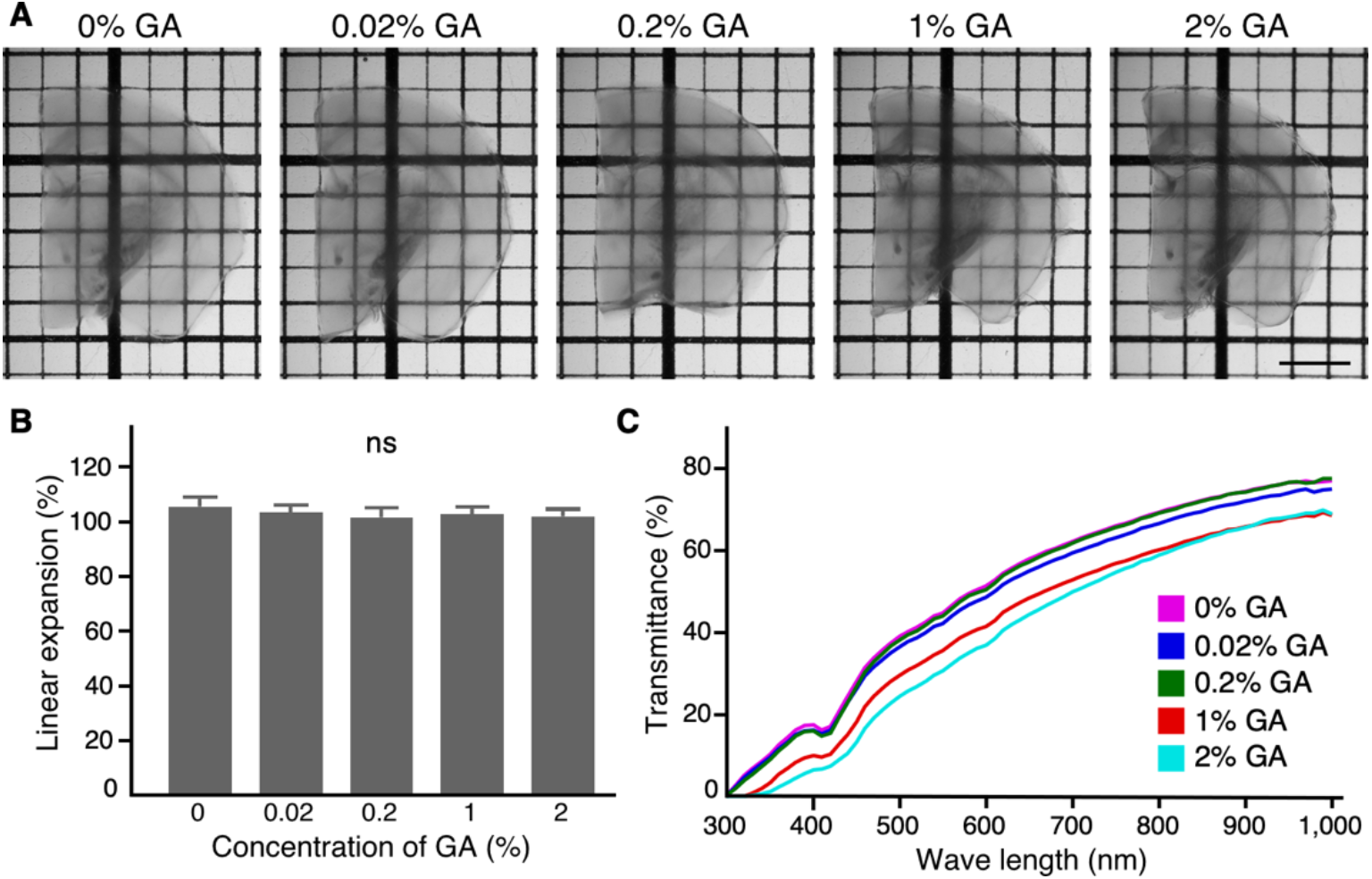
Sca*l*eSF clears brain slices fixed with GA. (**A**) Transmission images of Sca*l*eSF-treated mouse brain slices fixed with 4% PFA or 4% PFA containing GA (0.02, 0.2, 1 or 2%). The thickness of brain slices and the grid interval are 1 mm. (**B**) Change in size of brain slices after Sca*l*eSF treatment (n = 8, GA 0%; n = 8, GA 0.02%; n = 8, GA 0.2%; n = 8, GA 1%; n = 7, GA 2%; n = 4 mice for each condition; *F_4_,34 =* 1.975, *P =* 0.121, one-way ANOVA). Error bars represent SDs. (**C**) Transmission curves of Sca*l*eSF-treated mouse brain slices fixed with 4% PFA or 4% PFA containing GA (0.02, 0.2, 1, or 2%) (n = 3 brain hemispheres each). Scale bar: 2 mm.

**Fig. 3.**
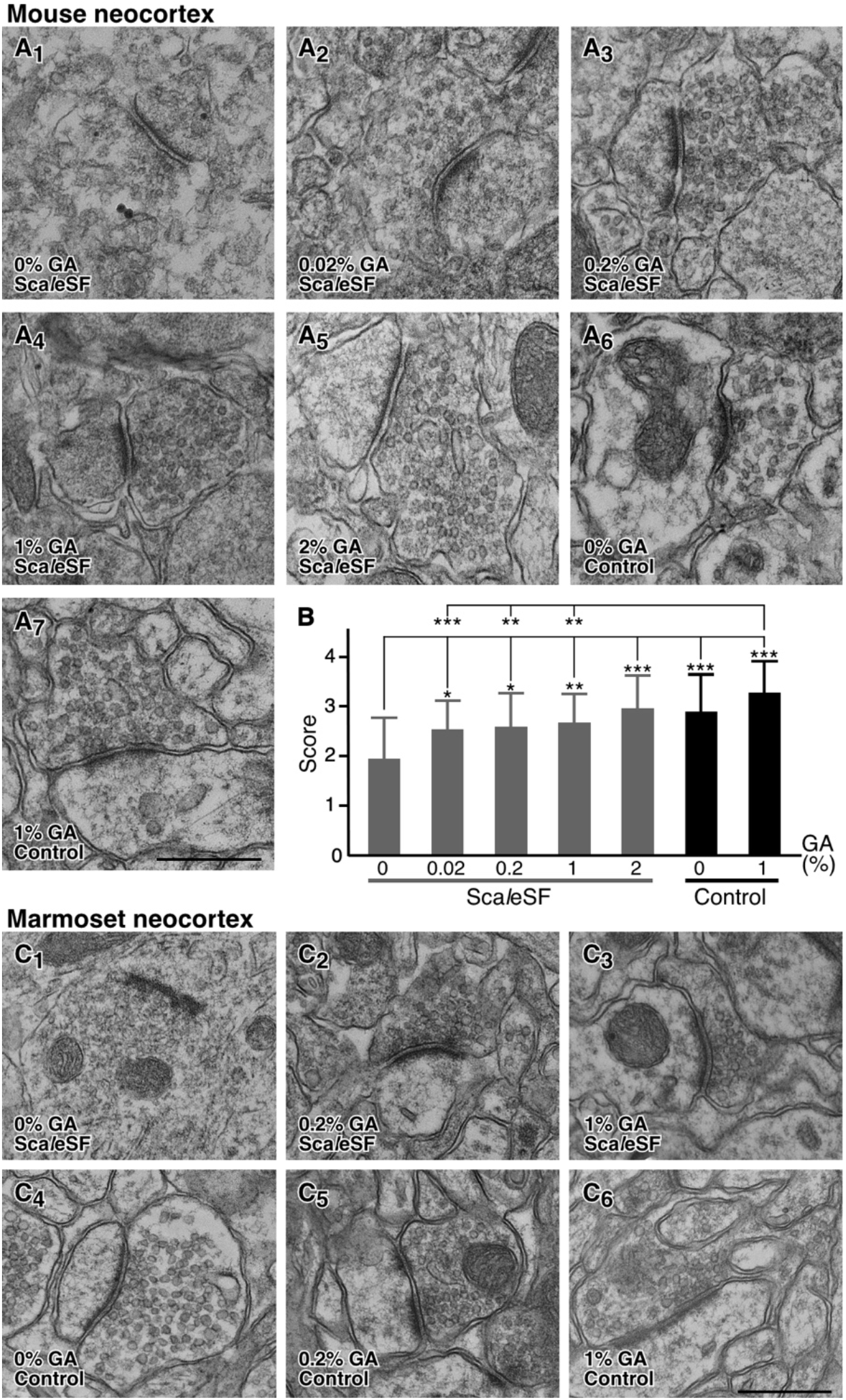
GA preserves ultrastructure in both mouse and marmoset brain slices cleared with Sca*l*eSF. (**A**) TEM images of mouse cerebral cortex cleared with Sca*l*eSF (A_1_ to A_5_) or stored in PBS(–) (A_6_, A_7_). Mice were fixed with 4% PFA (A_1_, A_6_) or 4% PFA containing GA (0.02%, A_2_; 0.2%, A_3_; 1%, A_4_, A_7_ or 2%, A_5_). (**B**) Scoring of membrane continuity of presynaptic terminals for each condition in (A). Over 90%, 50–90%, 10–50%, and less than 10% membrane continuity of presynaptic terminals are scored as 4, 3, 2, and 1, respectively (n = 31 synapses, GA 0%, Sca*l*eSF; n = 52 synapses, GA 0.02%, Sca*l*eSF; n = 33 synapses, GA 0.2%, Sca*l*eSF; n = 34 synapses, GA 1%, Sca*l*eSF; n = 31 synapses, GA 2%, Sca*l*eSF; n = 32 synapses, GA 0%, Control; n = 31 synapses, GA 1%, Control; n = 3 mice for each condition; *H* = 52.44, *df* = 6, *P =* 1.52 × 10-9 Kruskal–Wallis test; * *P <* 0.05, ** *P <* 0.01, *** *P <* 0.001; Steel–Dwass post hoc test). (**C**) TEM images of the cerebral cortex of marmosets. Ultrathin sections were prepared from brain slices cleared with Sca*l*eSF (C1 to C3) or stored in PBS(–) (C_4_ to C_6_). Marmosets were fixed with 4% PFA (C_1_, C_4_), 4% PFA containing 0.2% (C_2_, C_5_), or 1% GA (C_3_, C_6_) (n = 4 marmosets). Scale bars: 500 nm.

#### APEX2/BT-GO reaction that enables the correlated imaging of a fluorescent protein and an osmiophilic polymer in optically cleared tissues

For efficient successive LM/EM imaging in cleared tissues, we designed a genetically encoded probe for correlative light and electron microscopy (CLEM) by fusing EGFP in tandem with an engineered ascorbate peroxidase, APEX2 (EGFP-APEX2) (*27*). APEX2 catalyzes the polymerization and local deposition of 3,3-diaminobenzidine (DAB) in the presence of hydrogen peroxidase, which subsequently recruits electron-dense osmium to produce EM contrast. Importantly, APEX2 retains its peroxidase activity even after fixation with GA (*27–29*). We used a single adeno-associated virus (AAV) vector Tet-Off platform, AAV-SynTetOff (*30*), for high-level and neuronal expression of the CLEM probe (AAV2/1-SynTetOff-EGFP-APEX2-BGHpA) (Fig. 4A). We tested the feasibility of the vector by stereotactic injection into the mouse primary sensory cortex (S1). Seven to ten days after the injection, 1-mm-thick slices were prepared from the mouse brains and cleared with Sca*l*eSF. Tissue sections were cut perpendicularly to the deSca*l*ed slices (resectioning) and developed in the DAB-Ni^2+^ solution (Fig. 4B). Unexpectedly, DAB-Ni^2+^ labeling by APEX2 was much less sensitive than EGFP fluorescence-based detection in Sca*l*eSF-treated sections, hampering the correlated fluorescent and bright field imaging (Fig. 4C). We reasoned that clearing with Sca*l*eSF likely accounts for the lower sensitivity of APEX2, because DAB-Ni^2+^ labeling with APEX2 correlated well with EGFP fluorescence in untreated sections (Fig. 4D). To resolve this problem, we designed an experimental procedure in which biotin molecules are deposited with tyramide signal amplification (TSA) reaction using its peroxidase activity of APEX2 (APEX2/BT-GO reaction) prior to Sca*l*eSF treatment and then re-sections prepared from the cleared slices are processed for ABC/DAB-Ni^2+^ visualization (Fig. 4E). APEX2/BT-GO reaction gave remarkably strong DAB-Ni^2+^ labeling even after Sca*l*eSF treatment (Fig. 4F). DAB-Ni^2+^ labeling with APEX2/BT-GO reaction was comparable to or even more sensitive than EGFP fluorescence in Sca*l*eSF-treated sections (compare Fig. 4F_1_ with F_2_). Remarkably, we further observed DAB-Ni^2+^ labeling in fine subcellular structures such as axons, dendrites, and dendritic spines (Fig. 4F_3_ and F_4_). Thus, APEX2/BT-GO reaction combined with high-level gene transduction by the AAV-SynTetOff platform permits the correlated imaging of a fluorescent protein and an osmiophilic polymer in brain slices cleared with Sca*l*eSF.

**Fig. 4.**
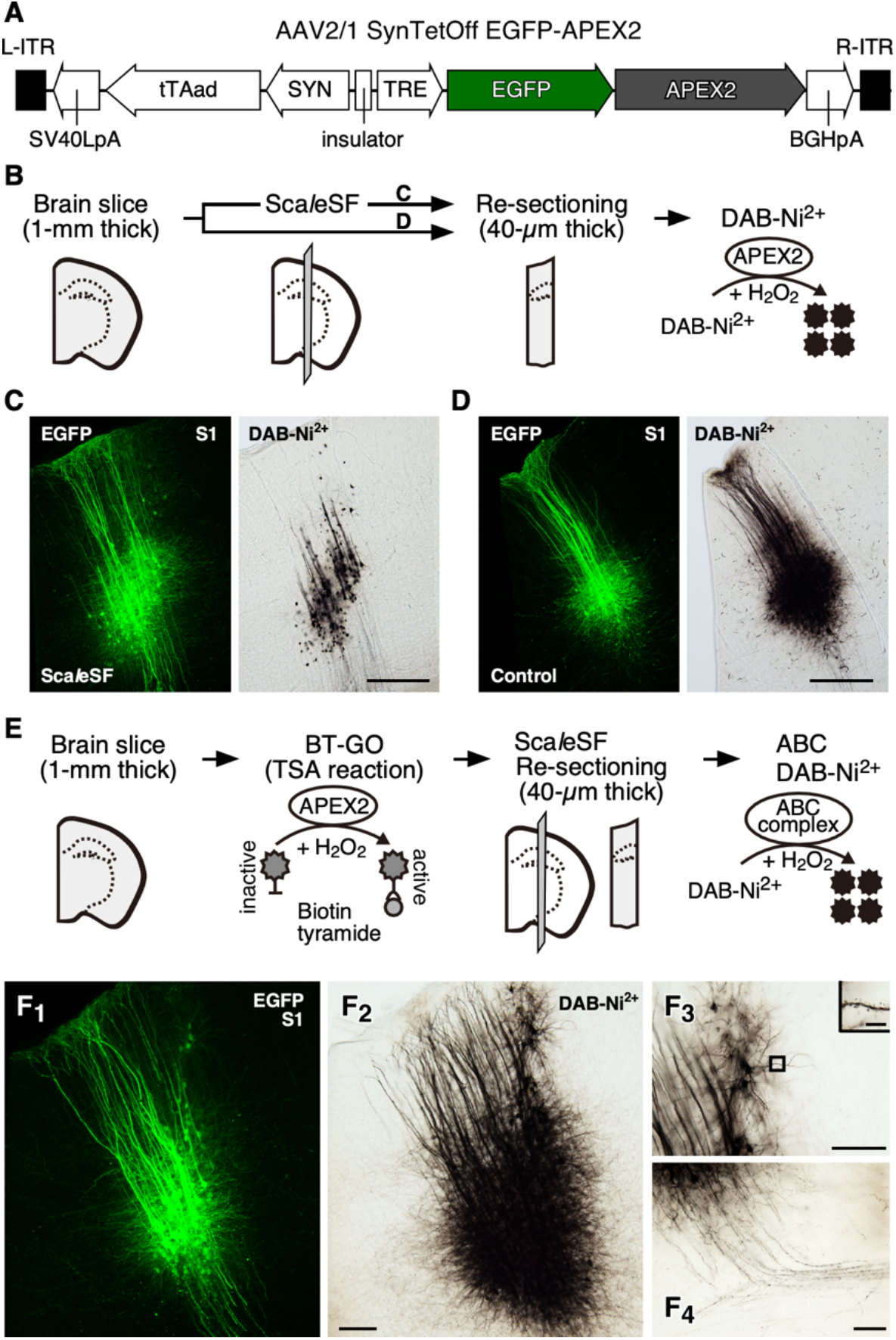
APEX2/BT-GO reaction enables correlated imaging of EGFP and DAB-Ni^2+^ polymers after Sca*l*eSF treatment. (**A**) AAV2/1-SynTetOff-EGFP-APEX2-BGHpA vector. (**B**) A schematic diagram of DAB-Ni^2+^ labeling with APEX2. (**C**, **D**) DAB-Ni^2+^ labeling with APEX2 in mouse brain sections prepared from brain slices cleared with Sca*l*eSF (C, n = 6 injection sites from 3 mice) or stored in PBS(–) (D, n = 7 injection sites from 4 mice). Correlated fluorescent (left) and bright-field (right) images in neurons infected with the AAV vector. After imaging with CLSM, sections are developed in DABNiNi2+ solution. (**E**) A schematic diagram of APEX2/BT-GO reaction-mediated signal amplification. Prior to clearing brain slices with Sca*l*eSF, biotin molecules are deposited with TSA reaction using its peroxidase activity of APEX2. The cleared slices are cut into 40-μm-thick sections and the sections are processed for ABC/DAB-Ni^2+^ visualization. (**F**) Correlated fluorescent (F_1_) and bright-field (F_2_) images in a section of a mouse cerebral cortex processed with APEX2/BT-GO reaction-mediated signal amplification (n = 7 injection sites from 4 mice). High magnification images of dendrites (F_3_), dendritic spines (the inset in F_3_), and axon fibers (F_4_) in the bright-field image are also shown. BGHpA: polyadenylation signal derived from the bovine growth hormone gene, ITR: inverted terminal repeat, SV40LpA: polyadenylation signal of Simian virus 40 late, SYN: human synapsin I promoter, TRE: tetracycline-responsive element, tTAad: an improved version of a tetracycline-controlled transactivator. Scale bars: 500 μm in (C, D), 100 μm in (F_2_), 50 μm in (F_3_, F_4_), and 5 μm in (the inset in F_3_).

#### Multi-scale LM/EM neuronal imaging in rodent and primate brains

By combining the aforementioned techniques, we implemented the multi-scale LM/EM neuronal imaging of three brain-wide circuits of two different species, mouse striatofugal, mouse callosal, and marmoset corticostriatal projection systems.

The caudate-putamen (CPu) is the primary input structure of the basal ganglia (*31*); it receives glutamatergic afferents from the cerebral cortex and thalamus, and sends GABAergic efferents to the external segment of the globus pallidus (GPe), entopeduncular nucleus (EP), and substantia nigra (SN). The striatofugal projection system was thus used as a model to test the feasibility of the imaging. The workflows for the multi-scale LM/EM neuronal imaging of murine striatal circuitry are presented in Fig. 5A. Four weeks after the injection of the AAV2/1-SynTetOff-EGFP-APEX2-BGHpA vector into the mouse CPu, the brains were fixed with 4% PFA containing 0.2% GA to improve ultrastructural preservation. Parasagittal slices (1-mm thick) were prepared from the brains, and biotin molecules were deposited with APEX2/BT-GO reaction. The slices were cleared with Sca*l*eSF, and then macroscopic and mesoscopic neural circuit mapping was conducted by CLSM (Fig. 5B). After perpendicular re-sectioning of the imaged slices (dotted lines in Fig. 5B), high-resolution image stacks were collected to document the detailed morphologies of the labeled neurons (Fig. 5C_1_–C_3_ and D_1_–D_3_). The imaged re-sections were processed for ABC/DAB-Ni^2+^ reaction using the deposited biotin molecules by APEX2/BT-GO reaction and embedded in an epoxy resin (Fig. 5C_4_ and D_4_). Ultrathin sections were prepared from the re-sections and imaged with TEM at a nanometer resolution (Fig. 5C_5_, C_6_, D_5_, and D_6_).

**Fig. 5.**
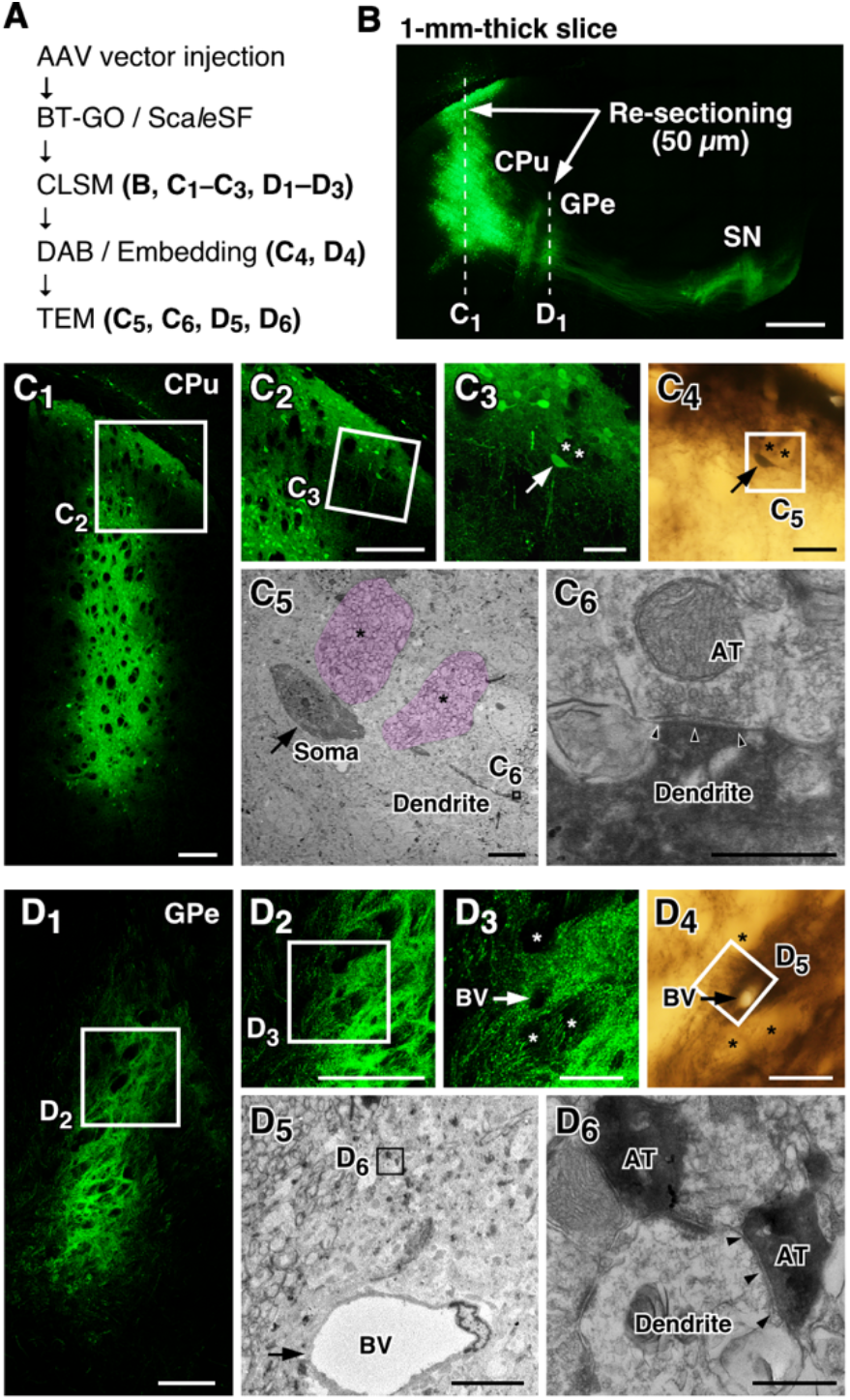
Multi-scale LM/EM neuronal imaging of mouse CPu neurons. (**A**) The procedure of multi-scale LM/EM neuronal imaging of mouse CPu neurons. (**B**) A maximum intensity projection image of a 1-mm-thick parasagittal brain slice cleared with Sca*l*eSF. AAV2/1-SynTetOff-EGFP-APEX2-BGHpA vector is injected into the CPu. Sections of 50-μm thickness are cut along dotted lines. (**C**, **D**) Correlated fluorescent (C_1_ to C_3_, D_1_ to D_3_), bright-field (C_4_, D_4_) and TEM images (C_5_, C_6_, D_5_, D_6_) at the level of CPu (C) and GPe (D). (**C_1_**, **D_1_**) CLSM imaging. (**C_2_**, **C_3_**, **D_2_**, **D_3_**) Enlarged views of rectangles in (C_1_), (C_2_), (D_1_), and (D_2_), respectively. (**C_4_**, **D_4_**) DAB-Ni^2+^ labeling with APEX2/BT-GO reaction. (**C_5_**, **D_5_**) A TEM image of the rectangle in (C_4_) and (D_4_). (**C_6_**, **D_6_**) A high magnification image of the rectangle in (C_5_) and (D_5_). A neuron indicated by arrows in (C_3_, C_4_) is targeted. Asterisks in (C_3_ to C_5_) and (D_3_, D_4_) indicate the same bundles of axonal fibers in (C) and (D), respectively. Arrows in (D_3_ to D_5_) indicate the identical blood vessel. Arrowheads in (C_6_, D_6_) indicate postsynaptic membranes. AT, axon terminal; BV, blood vessel. Scale bars: 500 μm in (B), 200 μm in (C_1_, C_2_, D_1_, D_2_), 50 μm in (C_3_, C_4_, D_3_, D_4_), 10 μm in (C_5_, D_5_), and 500 nm in (C_6_, D_6_).

We first performed the multi-scale LM/EM neuronal imaging of a synaptic input to a striatal neuron and a synaptic output to the GPe (Fig. 5). CLSM imaging in a Sca*l*eSF-treated brain slice clearly visualized the striatofugal projection system: EGFP-labeled fibers arising from the CPu extended caudally to the brainstem, forming dense terminal fields in the GPe and SN (Fig. 5B). We targeted a neuron in the dorsal CPu on the input side (Fig. 5C_1_–C_4_) and succeeded in performing EM imaging of the synaptic ultrastructure of the targeted dendrite (Fig. 5C_5_ and C_6_). The striatopallidal pathway, GABAergic inhibitory connections between the CPu and GPe, was mapped in the Sca*l*eSFtreated slice (Fig. 5B). A brain section from the imaged slice showed varicose axon arborization (Fig. 5D_1_ to D_3_) of the labeled neurons in the GPe. Following ABC/DAB-Ni^2+^ reaction (Fig. 5D_4_), axon terminals filled with the dark DAB precipitates were imaged with TEM (Fig. 5D_5_ and D_6_). We observed a symmetric synapse, which is characterized by the absence of postsynaptic densities (PSDs) and the narrow synaptic cleft, on a dendrite of the GPe neuron (Fig. 5D_6_). We then performed the multi-scale LM/EM neuronal imaging of striatonigral fibers on another cleared slice (Fig. S4). Myelin is a protein-lipid bilayer sheath that extends from oligodendrocytes and Schwann cells. Although an immunofluorescence study shows the unmyelinated character of striatonigral fibers (*32*), there is no direct evidence that striatonigral fibers are unmyelinated by EM observation. We thus applied our multi-scale LM/EM neuronal imaging to examine whether striatonigral fibers are indeed unmyelinated. Brain sections at the level of medial forebrain bundle (MFB) were prepared from the imaged slice and processed for successive LM/EM imaging (Fig. S4B and C). Targeting axonal bundles near the optic tract (OT) (Fig. S4C_1_ and C_2_), we found that almost all of the darkly stained axons were unmyelinated (Fig. S4C_3_ and C_4_). We also applied the multi-scale neuronal imaging to GABAergic inhibitory synapses between striatal projection neurons and SN neurons (Fig. S4B and D). Beginning with the mapping of the striatonigral projection in the cleared slice (Fig. S4B), varicose axon arborization was visualized in a re-section at the level of SN (Fig. S4D_1_), and a DAB-labeled axon terminal forming a symmetric synapse with a dendritic process was successively imaged (Fig. S4D_2_–D_4_).

Multi-scale LM/EM neuronal imaging should be effective in large-brained animals such as primates. The marmoset is becoming increasingly popular as a model organism in neuroscience research because of its social cognitive abilities and amenability to genetic manipulation (*33, 34*). We demonstrated the applicability of our multi-scale neuronal imaging in marmoset brains (Fig. 6). The AAV2/1-SynTetOff-EGFP-APEX2-BGHpA vector was injected into multiple neocortical sites of marmosets, and the brains were fixed with 4% PFA containing 0.2% GA. We identified clusters of neuronal elements visualized by EGFP-APEX2 expression in a macroscopic whole-brain image (Fig. 6B). The brains were then cut into 1-mm-thick coronal slices and those containing injection sites were cleared with Sca*l*eSF (Fig. 6C). Neural circuit mapping with CLSM clearly visualized the corticostriatal projection in the cleared slice: EGFP-labeled axons arising from the S1 extended subcortically and formed a dense terminal field in the putamen (Fig. 6D–F). After deSca*l*ing with PBS, the imaged slice was cut into sections for subcellular imaging with CLSM (Fig. 6G_1_ and H_1_).High-resolution image stacks in the re-sections documented the detailed morphologies of labeled neurons: pyramidal-shaped somata, apical and basal dendrites emanating from somata, and axonal projections extending basally and horizontally in the S1 (Fig. 6G_1_–G_3_), and axon terminal arborization and axonal boutons in the putamen (Fig. 6H_1_–H_3_). Of these structures, we targeted a basal dendrite of a pyramidal neuron on the input side (arrows in Fig. 6G_3_) and a corticostriatal axonal bouton on the output side (arrows in Fig. 6H_3_) for subsequent EM imaging. Following ABC/DAB-Ni^2+^ reaction and resin embedding (Fig. 6G_4_ and H_4_), ultrathin sections were prepared from the re-sections and further processed for EM (Fig. 6G_5_, G_6_, H_5_, and H_6_). We observed an asymmetric synapse, which typically mediates glutamatergic neurotransmission, on the targeted dendrite filled with electron-dense DAB precipitates (Fig. 6G_6_), as well as an asymmetric synapse between a corticostriatal axon terminal and a striatal dendrite (Fig. 6H_6_). Given macroscopic imaging of centimeter-sized marmoset brains (3 cm length and 2 cm width; Fig. 6B) and TEM imaging of synapses with nanometer resolution (1.2 nm/pixel; Fig. 6G_6_ and H_6_), we succeeded in multi-scale LM/EM neuronal imaging with over seven orders of magnitude.

**Fig. 6.**
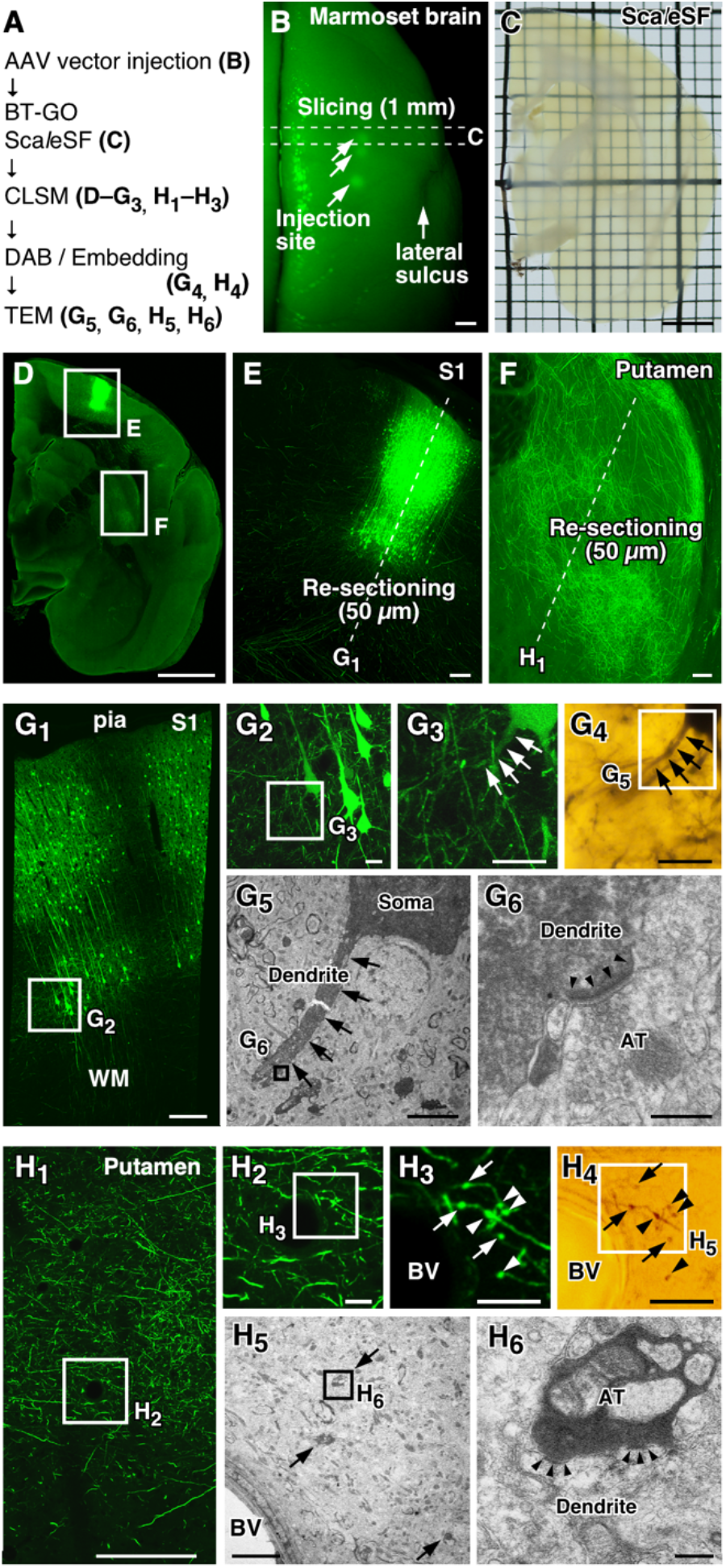
Multi-scale LM/EM neuronal imaging of cortical neurons in a marmoset. (**A**) The procedure of multi-scale LM/EM neuronal imaging. (**B**) EGFP fluorescence (arrowheads) in a marmoset brain six weeks after injection of the AAV2/1-SynTetOff-EGFP-APEX2-BGHpA vector. A 1-mm-thick slice is cut along dotted lines. (**C**) A transmission image of the slice cleared with Sca*l*eSF. (**D–F**) Maximum intensity projection images of the cleared slice. (**E**, **F**) Enlarged views of the S1 (E) and putamen (F). Sections of 50 μm thickness are cut along dotted lines. (**G**, **H**) Correlated fluorescent (G_1_ to G_3_, H_1_ to H_3_), bright-field (G_4_, H_4_), and TEM images (G_5_, G_6_, H_5_, H_6_) in the S1 (G) and putamen (H). (**G_1_, H_1_**) CLSM imaging. (**G_2_, G_3_, H_2_, H_3_**) Enlarged views of the rectangles in (G_1_), (G_2_), (H_1_), and (H_2_), respectively. (**G_4_, H_4_**) DAB-Ni^2+^ labeling with APEX2/BT-GO reaction. (**G_5_, H_5_**) TEM images of the rectangle in (G_4_) and (H_4_). (**G_6_, H_6_**) A high magnification image of the rectangle in (G_5_) and (H_5_). A synaptic structure (arrowheads in G_6_) in a dendrite (arrows in G_3_ to G_5_) of a pyramidal neuron is targeted in (H) and synaptic structures (arrowheads in H_6_) between a cortical axon and a putamen dendrite are targeted in (G). Arrows in (H_3_ to H_5_) and arrowheads in (H_3_, H_4_) indicate the same presynaptic terminals. BV: blood vessel. Scale bars: 3 mm in (B to D), 200 μm in (E, F, G_1_, H_1_), 20 μm in (G_2_ to G_4_, H_2_ to H_4_), 5 μm in (G_5_, H_5_), and 300 nm in (G_6_, H_6_).

Our multi-scale LM/EM imaging allows for high-speed LM imaging of substantial tissue volume at high-resolution and subsequent EM observation of targeted structures, facilitating the capture of scarce structures with nanoscale resolution. Callosal projection neurons are a heterogenous population of neocortical projection neurons that interconnect the two hemispheres of the cerebral cortex (*35*). Notably, callosal inputs onto GABAergic neocortical interneurons are scant: the vast majority of callosal terminals synapses onto dendritic spines, likely those of excitatory pyramidal neurons, while the remainder synapses onto dendritic shafts of spiny and aspiny neurons in mice (*36, 37*). We therefore chose callosal synaptic inputs onto a neocortical GABAergic interneuron subtype, PV neocortical interneurons, in mice as scarce structures with nanoscale resolution and tracked them in a targeted way across multiple spatial scales by successive LM/EM imaging (Fig. 7). The AAV2/1-SynTetOff-EGFP-APEX2-BGHpA vector was injected into the primary motor cortex (M1) and the AAV2/1-SynTetOff-FLEX-mScarlet-BGHpA was injected into the contralateral M1 of a *PV^+/Cre^* mouse to label callosal axons with EGFP and PV neocortical interneurons with mScarlet (Fig. 7A and B). CLSM imaging in a Sca*l*eSF-treated brain slice mapped the callosal projection system: EGFP-labeled axons arising from the M1 passed through the corpus callosum and projected to the homotopic contralateral cortex, where mScarlet-labeled PV interneurons were located (Fig. 7C and D). We screened a large number of (> 3000) serial *xy* images (121 × 121 μm square) at different *z* positions in a thick brain slice (1-mm thickness) cleared with Sca*l*eSF and identified an apposition between a callosal axon terminal and a dendrite of PV neocortical interneuron (Fig. 7E, arrowhead). After re-sectioning the imaged slices parallel to the *xy* plane (parallel re-sectioning) followed by counterstaining with DAPI (4’,6-diamidino-2-phenylindole), the possible synaptic contact was validated with high-resolution imaging with CLSM (Fig. 7F, arrowheads). Following ABC/DAB-Ni^2+^ reaction and resin embedding, the re-section was subjected to FIB-SEM imaging (Fig. 7G–I and Movie S1). The CLSM image in the slice exactly matched the FIB-SEM tomogram (compare Fig. 7E with H): mScarlet fluorescence corresponded with the SEM profile of membrane structure, and EGFP fluorescence correlated well with the DAB-Ni2+ precipitates. Correlation of CLSM in the Sca*l*eSF-treated brain slice, CLSM in the re-section, and FIB-SEM datasets demonstrated the preservation of structural integrity throughout successive LM/EM imaging (Fig. 7C–I). The axodendritic apposition between a callosal axon and a PV neocortical interneuron (Fig. 7E and F, arrowheads) actually formed a synaptic contact: we observed an asymmetric synaptic specialization, which is characterized by the existence of PSD, at the apposition between the axon terminal filled with electron-dense DAB precipitates and the dendrite in a FIB-SEM tomogram (Fig. 7I).

**Fig. 7.**
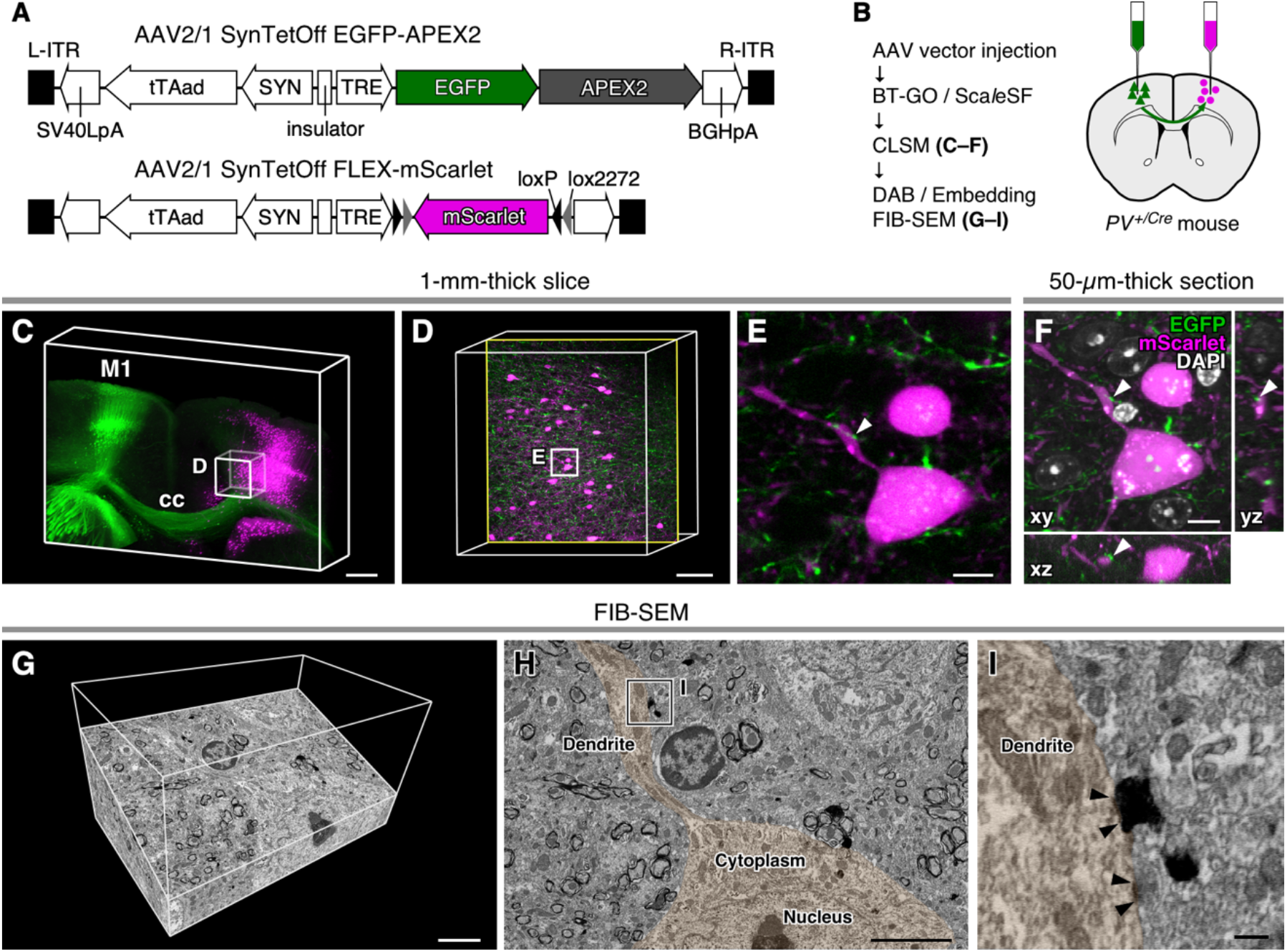
Multi-scale LM/EM neuronal imaging of a mouse callosal synaptic input onto PV neocortical interneurons. (**A**) AAV2/1-SynTetOff-EGFP-APEX2-BGHpA vector and AAV2/1-SynTetOff-FLEX-mScarlet-BGHpA vector. (**B**) The procedure of multi-scale LM/EM neuronal imaging of mouse callosal synaptic inputs onto PV neocortical interneurons. **(C–E)** CLSM imaging of a 1-mm-thick brain slice. (**C**) Three-dimensional volume rendering of the M1 of a *PV^+/Cre^* mouse injected with the AAV vectors. (**D**) An enlarged and high-resolution image of the box in (C). The image is 90° rotated in a counterclockwise direction with respect to (C). An optical section with an axodendritic apposition in (E) is shown. (**E**) A higher magnification image of the rectangle in (D). (**F**) CLSM imaging in a re-section. A 50-μm-thick section was cut parallel to the *xy* plane from the slice imaged in (C to E). Orthogonal views of the *xz* plane (bottom) and the *yz* (right) are also shown. Arrowheads in (E, F) indicate the same axodendritic apposition. (**G–I)** FIB-SEM tomography of the axodendritic apposition. (**G**) A three-dimensional volume rendering image. (**H**) An oblique slice view. (**I)** An enlarged view of the rectangle in (H). Arrowheads in (I) indicate the PSD. The profiles of postsynaptic dendrite and soma of the targeted axodendritic apposition (arrowheads in E, F) are pseudocolored in (H, I) for clarity. BGHpA: polyadenylation signal derived from the bovine growth hormone gene, cc: corpus callosum, ITR: inverted terminal repeat, SV40LpA: polyadenylation signal of Simian virus 40 late, SYN: human synapsin I promoter, TRE: tetracycline-responsive element, tTAad: an improved version of a tetracycline-controlled transactivator. Scale bars: 500 μm in (C), 100 μm in (D), 10 μm in (E, F), 5 μm in (G, H), and 500 nm in (I).

### Discussion

The imaging scale required for deciphering brain-wide connectivity in the mammalian brain exceeds several orders of magnitude (*3*). We overcame the technical challenges associated with this requirement by coupling a tissue clearing method with successive LM/EM imaging. Our multi-scale LM/EM neuronal imaging enables brain-wide connectomic analysis by the simultaneous interrogation of their neural circuit structures with LM and synaptic connectivity with EM. The feasibility of the multi-scale neuronal imaging was demonstrated in the mouse striatofugal projection system. Beginning with mapping the projection in cleared brain tissues, we anterogradely imaged the detailed morphologies of labeled neurons with CLSM and targeted nanoscopic structures such as synapses and myelin sheaths (Fig. 5 and Fig. S4). As demonstrated by the application to marmoset brains (Fig. 6), our multi-scale imaging should be effective in connectomic analysis of large-brained animals. Our multi-scale LM/EM imaging that is featured with high-speed and high-resolution LM imaging followed by subsequent EM imaging at a nanometer resolution allowed us to capture scarce synaptic contacts, callosal inputs onto PV neocortical interneurons in mice (Fig. 7). Our multi-scale LM/EM imaging can complement current comprehensive connectomic analysis (*4, 5*). While current comprehensive approaches with EM alone are mainly applied to small pieces of brain tissues (*6–9*), the present imaging modality makes it possible to describe synaptic connectivity of brain-wide circuits by integrating seamlessly structural information with different spatial scales in a reasonable amount of time without specialized equipment.

Sca*l*eSF, a rapid, isometric, and GA-resistant tissue clearing technique, facilitated multi-scale LM/EM neuronal imaging. Multi-scale LM/EM imaging requires a tissue clearing method that allows for the preservation of ultrastructure and fluorescence signals. However, most tissue clearing methods, especially protocols featuring high clearing capabilities, aggressively remove lipids and pigments for extensive tissue clarification (*15, 16*), compromising ultrastructural preservation (*22 38*) (but see ref. (*39*)). Compared to solvent and hydrogel-based tissue clearing methods, aqueous tissue clearing methods surpass in preserving fluorescence signals and tissue integrity (*15, 16*). Although aqueous tissue clearing methods containing minimal lipid-extracting detergents have been reported (*15, 16*), none are suitable for use with our multi-scale LM/EM imaging, i.e., isometricity, resistance against GA, and a clearing capability for 1-mm-thick brain slices. The 1-mm thickness of the brain slices used in this study is satisfactory enough to recover all of dendritic arbors and inhibitory interneuron axonal arbors of the rodent and carnivore cerebral cortex in their entirely (*40, 41*), providing rich structural information on neural circuit architecture.

Although Sca*l*eSF achieved a high level of preservation of ultrastructure and fluorescence signals (Fig. 1–3), two challenges remain in the clearing protocol. First is the advanced preservation of ultrastructure: a slight but statistically significant degradation of the ultrastructure in brain slices cleared with Sca*l*eSF (Fig. 3B) leaves a room for further improvement. Second is the scaling of the clearing protocol: Sca*l*eSF was developed for clearing brain slices, not for a whole brain. Although 1-mm-thick brain slices provide good knowledge of dendritic and local axonal arbors, information about long-range projections is fragmentary and incomplete in the slice (*12, 13, 41–43*). The direct perfusion of clearing reagents that enhances clearing capability (*44–46*) might permit whole-brain clearing accompanied with preserved ultrastructure and fluorescence signals.

Successive LM/EM imaging can be performed efficiently with fluorescent and electron-dense genetically encoded CLEM probes. LM/EM dual labeling with a single protein enables the unambiguous correlation of LM and EM datasets. Although the correlation can be achieved by endogenous and artificial landmarks, these techniques require additional labeling for endogenous landmarks and/or specialized equipment (*47–49*). Genetically encoded CLEM probes for our multiscale LM/EM neuronal imaging should be stable in cleared samples. APEX2 retains its peroxidase activity even upon fixation with GA (*27–29*), rendering APEX2 fusion constructs with fluorescent proteins as good candidates for the CLEM probes. However, we found that its peroxidase activity of APEX2 was unexpectedly low after clearing with Sca*l*eSF (Fig. 4C and D). Hence, we introduced APEX2/BT-GO reaction prior to the clearing treatment to deposit biotin molecules with TSA reaction using its peroxidase activity of APEX2 (Fig. 4E). APEX2/BT-GO reaction provided remarkably strong DAB-Ni^2+^ labeling while maintaining EGFP fluorescence (Fig. 4F) that achieved LM/EM dual labeling in brain slices even cleared with Sca*l*eSF. Despite the potent LM/EM dual labeling with APEX2 BT-GO reaction, the reaction itself and permeabilization with a lipid-extracting detergent can potentially damage cellular ultrastructure. Our LM/EM dual labeling coupling a genetically encoded CLEM probe, EGFP-APEX2, with APEX2/BT-GO reaction gave strong EM contrast throughout the cytoplasm (Fig. 5–7 and Fig. S4). Although cytoplasmic labeling with DAB facilitates the identification of targeted structures, the labeling may interfere with the visualization of ultrastructural features of synapses such as active zone, synaptic vesicle morphologies, and PSDs. Peroxidase constructs targeted to subcellular compartments would make it possible to visualize ultrastructural properties of synaptic arrangements as well as multiplexed labeling in EM (*50*).

The simultaneous interrogation of molecular and structural information is required for the advancement of connectomic analysis. However, molecular information is often lost in connectomic analysis with EM alone, and LM lacks nanoscale resolution necessary to identify a single synapse. Our multi-scale LM/EM neuronal imaging overcomes the deficiency of both analyses. Sca*l*e technologies achieve stable tissue preservation for immunohistochemical labeling on re-sections prepared from deSca*l*ed tissues (K. Y. and H. H., unpublished observations) (*22, 26*) and can thus be used to collect both molecular and structural information. Furthermore, our labeling approach with genetically encoded probes can be applied to a library of Cre driver lines, providing us with a genetic handle on studying neural circuit structure and synaptic connectivity of specific neuronal types. Indeed, we identified and tracked mouse callosal inputs onto a neocortical GABAergic interneuron subtype, PV neocortical interneurons, by injecting a flexed AAV vector coding for mScarlet into *PV^+/Cre^* mouse brains (Fig. 7). The high-level preservation of fluorescent signals and ultrastructure in Sca*l*eSF-treated brain slices (Fig. 1–3) is amenable to *post hoc* molecular mapping with high accuracy on re-sections, such as array tomography (*51*) and super-resolution imaging (*17*).

In summary, we developed and validated multi-scale LM/EM neuronal imaging for connectomic analysis of neuronal circuits spanning the mammalian brain. Our imaging modality will significantly advance the understanding of brain-wide connectivity by expanding the scales of objects.

### Materials and Methods

#### Animals

All animal experiments involving animal care, surgery, and sample preparation were approved by the Institutional Animal Care and Use Committees of Osaka University (Approval No. 300150), Juntendo University (Approval No. 2020087, 2020088), and Kyoto University (Approval No. Med Kyo 20031) and conducted in accordance with Fundamental Guidelines for Proper Conduct of Animal Experiments by the Science Council of Japan (2006). All efforts were made to minimize animal suffering and the number of animals used.

Eight- to twelve-week-old male C57BL/6J (Nihon SLC), *PV^Cre^* heterozygous (*Pvalb^tm1(cre)Arbr^*, The Jackson Laboratory Stock No: 008069) (*52*), and PV/myristoylation-EGFP-low-density lipoprotein receptor C-terminal BAC transgenic mice (PV-FGL mice) (*24*) under specific pathogen-free (SPF) conditions were used. The mice were maintained under a 12/12 hr light/dark cycle (light: 08:00–20:00) with ad libitum access to food and water. Mouse genotypes were determined by polymerase chain reaction (PCR) analysis as described previously (*24*).

Four young adult (14–15 months old) male or female common marmosets (Callithrix jacchus; body weight, 280–400 g; bred either in CLEA Japan or in our laboratory) were housed in their home cages under a 14/10 hr light/dark cycle (light: 07:00–21:00). Each cage had a wooden perch, a food tray, and an automatic water dispenser. The animals were fed twice a day with solid food (CMS-1, CLEA Japan). Water was provided ad libitum.

#### Preparation of tissue slices

Mice were deeply anesthetized by an intraperitoneal injection of sodium pentobarbital (200 mg/kg; Somnopentyl, Kyoritsu Seiyaku) and perfused transcardially with 20 mL of 5 mM phosphate-buffered 0.9% saline (PBS; pH 7.4) at 4°C, followed by the same volume of 4% paraformaldehyde (PFA) (1.04005.1000, Merck Millipore) or 4% PFA containing various concentrations (0.02, 0.2, 1 or 2%) of glutaraldehyde (GA) (17003-92, Nacalai Tesque) in 0.1 M phosphate buffer (PB; pH 7.4) at 4°C. The brains of the animals were removed and postfixed in the same fixatives overnight at 4°C. After embedding in 4% agar (01028-85, Nacalai Tesque) in PBS, coronal or sagittal slices of 1-mm in thickness were cut with a vibrating tissue slicer (Linear PRO7N, Dosaka EM).

The maromsets were deeply anesthetized by an intramuscular injection of ketamine (60 mg/kg; Ketalar, Daiichi Sankyo Propharma) and an intraperitoneal injection of sodium pentobarbital (80 mg/kg). The fixation and preparation of the tissue slices of marmoset brains were the same as those used for the mice, except that their perfusion with 300 mL of 1.0 unit/mL heparin (224122458, Mochida Pharmaceutical) in PBS followed by the same volume of 4% PFA or 4% PFA containing GA (0.2% or 1%) in 0.1 M PB.

#### Tissue clearing

The schedule for tissue clearing with Sca*l*eSF is described in Fig. 1A. Brain slices were permeabilized with Sca*l*eS0 solution for 2 hr at 37°C, washed twice with PBS(–) (27575-31, Nacalai Tesque) for 15 min at 20–25°C, and cleared with Sca*l*eS4 solution for 8–12 hr at 37°C. We treated brain slices with Sca*l*eS4 solution for 12 hr in the data shown in this paper. The formula for Sca*l*eS0 solution was 20% (w/v) sorbitol (06286-55, Nacalai Tesque), 5% (w/v) glycerol (G9012, Sigma-Aldrich), 1 mM methyl-β-cyclodextrin (M1356, Tokyo Chemical Industry), 1 mM γ-cyclodextrin (037-10643, Wako Pure Chemical Industries), and 3% (v/v) Dimethyl Sulfoxide (DMSO) (13407-45, Nacalai Tesque) in PBS(–), and that for Sca*l*eS4 solution was 40% (w/v) sorbitol, 10% (w/v) glycerol, 4 M urea (35940-65, Nacalai Tesque), 0.2% (w/v) Triton X-100 (35501-15, Nacalai Tesque), and 25% (v/v) DMSO in distilled deionized water(*23*).

Optical clearing of brain slices with Sca*l*eSQ(0) was performed as described previously (*22*).

#### Observation and measurement of macroscopic structures

Transmission images of mouse brain slices were acquired with a stereomicroscope (M205C, Leica Microsystems) equipped with a 1× objective lens (PLANAPO, working distance [WD] = 65 mm, Leica Microsystems), a transmitted light base (TL RCI™, Leica Microsystems), and a digital single lens reflex camera (D7200, Nikon). Marmoset brain slices were placed on a LED tracing board (A4-500, Trytec) and imaged with the digital single lens reflex camera mounted on a copy stand (CS-A4 L18142, LPL). Fluorescence images of the marmoset brains were acquired with the stereomicroscope equipped with an external fluorescence light source (EL6000, Leica Microsystems), a GFP filter cube (excitation filter: 470 ± 20 nm, emission filter: 525 ± 25 nm, Leica Microsystems), and a cooled CCD camera (Rolera-XR, QImaging). Brain samples were placed on graph paper with a patterned background (ruled into 1-mm squares).

To assess tissue expansion or shrinkage caused by tissue clearing, brain-slice areas were measured with ImageJ (ver. 1.52v, National Institutes of Health) (*53*). Linear expansion values were determined based on the square root of the changes in brain-slice areas.

#### Transmission measurements

Light transmittance of brain slices was determined with a spectrofluorometer (Enspire 2300, Perkin Elmer). Coronal brain slices at the level of the S1 were used. Brain slices cleared with Sca*l*eSF or Sca*l*eSQ(0), or stored in PBS(–) were transferred onto UV transparent 96-well plates (655801, Greiner Bio-One) to measure absorbance of the tissues. The absorbance (A) was converted to percent transmittance (%T) using an equation derived from Lambert-Beer’s law: A = 2 – log10 (%T).

#### Imaging chamber and tissue mounting

A customizable 3D-printed imaging chamber that enabled the reliable mounting of cleared tissues was designed for imaging with CLSM (Fig. S1). The chamber consisted of the chamber frame, bottom coverslip, and microscope stage adaptors (Fig. S1A). The frames and adaptors were designed according to the size and thicknesses of brain slices and ordered to be printed from a rigid acrylic resin, AR-M2 (Keyence), using a 3D-printer (AGILISTA-3200, Keyence) by DMM.make (https://make.dmm.com). The frames were glued to the bottom coverslips (Matsunami). Optically cleared tissues were mounted on the coverslips and embedded in 1.5% Agarose (L03, TaKaRa Bio) in Sca*l*eS4D25(0) solution (Sca*l*eS4 gel) (*23*). Tissues were coverslipped and left at 4°C until the gel solidified. The imaging chambers were attached to the microscope stage adaptors to mount on microscope stages (Fig. S1B and C) or attached to petri dishes with Blu-Tak^®^ (Bostik) and immersed in Sca*l*eS4 solution (Fig. S1D).

#### Confocal laser scanning microscope

3D image stacks were acquired with a TCS SP8 CLSM (Leica Microsystems). A 16× multi-immersion objective lens (HC FLUOTAR 16x/0.60 IMM CORR VISIR, numerical aperture [NA] = 0.60, WD = 2.5 mm, Leica Microsystems) was used for imaging the optically cleared brain slices (1-mm thick). A 10× air (HCX PL APO 10x/0.40 CS, NA = 0.40, WD = 2.20 mm, Leica Microsystems), a 20× multi-immersion (HC PL APO 20x/0.75 IMM CORR CS2, NA = 0.75, WD = 0.68 mm, Leica Microsystems), a 25× water-immersion (HC FLUOTAR L 25x/0.95 W VISIR, NA = 0.95, WD = 2.50 mm, Leica Microsystems), and a 63× oil-immersion (HC PL APO 63x/1.40 Oil CS2, NA = 1.40, WD = 0.14 mm, Leica Microsystems) objective lenses were used for imaging the sections (40- or 50-μm thick). Sections were mounted with PBS or 75% glycerol in PBS (*29*). DAPI, EGFP, and mScarlet were excited by 405-, 488-, and 552-nm lasers, and their fluorescence was collected through 410–480, 495–525, and 560–700 nm emission prism windows, respectively.

#### Transmission electron microscopy

Sample preparation and imaging of the cleared brain slices with TEM were carried out as described previously (*22*), with minor modifications. Briefly, 1-mm-thick brain slices were cleared with Sca*l*eSF and embedded in Sca*l*eS4 gel for 24 hr or stored in PBS(–) at 4°C. 1-mm-cubes were excised from the brain slices with carbon steel blades (FA-10B, Feather). The cubes and re-sections (50-μm thick) prepared from 1-mm-thick slices were osmicated with 1% OsO4 (25746-06, Nacalai Tesque) in 0.1 M PB, dehydrated with a gradient series of ethanol (50, 70, 90, 99, and 100%) followed by propylene oxide (29223-55, Nacalai Tesque), and embedded in an Epon 812 mixture (a mixture of Luveak-812 [20829-05, Nacalai Tesque], Luveak-DDSA [14423-95, Nacalai Tesque], Luveak-MNA [14424-85, Nacalai Tesque], and Luveak-DMP-30 [14425-75, Nacalai Tesque]) or Durcupan (44610, Sigma-Aldrich). To test the accelerator Luveak-DMP-30 for the permeability of the Epon 812 mixture into the tissues, resin-polymerization was initiated after pre-incubation with an Epon 812 mixture that did not contain the accelerator (modified Epon method). After polymerization of the resin, ultrathin sections (70-nm thick) were cut with an ultramicrotome (Ultracut UCT, Leica Microsystems). The sections were stained with 1% uranyl acetate and 1% lead citrate, and were observed under a TEM (H-7650, Hitachi) at 80 kV. We acquired digital photographs of presynaptic axonal terminals, which contained synaptic vesicles and synapsed with dendritic structures, at a resolution of 1.2 nm/pixel.

To evaluate ultrastructural preservation, the plasma membrane of the presynaptic structures was outlined with a graphic software (CANVAS X DRAW, ACD systems). Membrane continuities of presynaptic structures of > 90%, 50–90%, 10–50%, and < 10% were scored as 4, 3, 2, and 1, respectively.

#### Scanning electron microscopy combined with focused ion beam

We also performed 3D imaging of synaptic structures by FIB-SEM technique as described previously (*54*), with minor modifications. In brief, brain sections (50-μm thick) were osmicated with 2% OsO4 in 0.1 M PB, counterstained with 1% uranyl acetate for 2 hr, and stained in lead aspartate solution at 60°C for 30 min. After dehydration with a gradient series of ethanol (60, 70, 80, 90, 99, and 100%) and propylene oxide, the sections were flat-embedded in the Epon 812 mixture. The regions that contained targeted structures were excised by carbon steel blades from the embedded sections, mounted on aluminium stubs, and examined with a FIB-SEM system (Crossbeam 540, Carl Zeiss Microscopy). Using the FIB of 30 kV and 3 nA, a surface layer of 10-nm thickness was milled at each sectioning. Following the removal of each layer, the freshly exposed surface was imaged with the SEM using the back-scattered electron detector at a magnification of 10 nm/pixel. The acceleration voltage of the imaging beam was 1.5 kV with a beam current of 1 nA and a dwell time of 13.6 μs/pixel.

#### AAV vector construction and production

pAAV2-SynTetOff-EGFP-APEX2-BGHpA was constructed as follows. The GFP sequence of pENTR1A-SV40LpA-tTAad-SYN-insulator-TRE-GFP-BGHpA (*30*) was replaced with a multiple cloning site, which contained BamHI-BglII-SalI restriction sites. The resultant entry vector pENTR1A-SV40LpA-tTAad-SYN-insulator-TRE-BBS-BGHpA, namely pENTR1A-SynTetOff-BBS-BGHpA, was reacted with pAAV2-DEST(f) (*30*) by homologous recombination with LR clonase II (11791020, Thermo Fisher Scientific) to generate pAAV2-SynTetOff-BBS-BGHpA. A DNA fragment encoding EGFP-APEX2 fusion protein was generated by overlapping PCR. A sequence coding for a peptide linker (Gly-Gly-Ser)2 was inserted between the two protein domains. The coding sequence of APEX2 was amplified from pcDNA3-Connexin43-GFP-APEX2 (#49385, Addgene) (*27*). The restricted products were inserted into pAAV2-SynTetOff-BBS through the BamHI/SalI sites, resulting in pAAV2-SynTetOff-EGFP-APEX2-BGHpA. For the construction of pAAV2-SynTetOff-FLEX-mScarlet-BGHpA, the coding sequence of mScarlet was amplified from pmScarlet_C1 (#85042, Addgene) (*55*). The amplified products were inserted into pBSIISK-hFLEX (*30*) through the PstI/EcoRI sites to generate pBSIISK-FLEX-mScarlet. The pBSIISK-FLEX-mScarlet was then digested with BamHI/SphI and ligated into the corresponding sites of pAAV2-SynTetOff-BBS, yielding pAAV2-SynTetOff-FLEX-mScarlet-BGHpA. The following primers were used for the PCR amplification: BamHI-kozak-EGFP: 5’-AAAAGGATCCGCCACCATGGTGAGCAAGGG-3’, EGFP-(GGS)2: 5’-GGAACCACCGGAACCACCCTTGTACAGCTCGTCCATGC-3’, (GGS)2-APEX2: 5’-GGTGGTTCCGGTGGTTCCGGAAAGTCTTACCCAACTGT-3’, SalI-APEX2: 5’-TTTTGTCGACTTAGGCATCAGCAAACCCAA-3’, Pstl-kozak-mScarlet: 5’-AAAACTGCAGATGGTGAGCAAGGGCGAGGC-3’, and mScarlet-stop-EcoRI: 5’-TTTTGAATTCTTACTTGTACAGCTCGTCCATGC-3’.

AAV vector particles were produced and purified as described previously (*30*). Briefly, pAAV2-SynTetOff-EGFP-APEX2-BGHpA or pAAV2-SynTetOff-FLEX-mScarlet-BGHpA and two helper plasmids were co-transfected into HEK293T cells (RCB2202, Riken) using polyethylenimine (23966, Polysciences). Virus particles were purified from the cell lysate or the cell lysate and supernatant by ultracentrifugation with OptiPrep (1114542, Axis-Shield) and concentrated by ultrafiltration with Amicon Ultra-15 (UFC905024, Merck Millipore). The infectious titer of the AAV vector (IFU/mL) was determined by quantitative PCR (qPCR) with HEK293T cells infected with the purified AAV vectors. The physical titer of the AAV vector (genome copies (gc)/mL) was measured by qPCR with the purified viral solutions. The solution was stored in aliquots at −80°C until use.

#### Virus injection

Virus injection into mouse brains was carried out as described previously, with some modifications (*56, 57*). Briefly, mice were deeply anesthetized with an intraperitoneal injection of medetomidine (0.3 mg/kg; Domitor, Zenoaq), midazolam (4 mg/kg; Dormicum, Astellas Pharma), and butorphanol (5 mg/kg; Vetorphal, Meiji Seika Pharma) and placed in a stereotaxic apparatus (SR50, Narishige). Subsequently, 0.2 μl of the viral solution (AAV2/1-SynTetOff-EGFP-APEX2-BGHpA: 1.32 × 1011 IFU/mL, AAV2/1-SynTetOff-FLEX-mScarlet-BGHpA: 1.8 × 1013 gc/mL) was pressure injected into the M1, S1, and CPu through a glass micropipette attached to Picospritzer III (Parker Hannifin). The injection coordinates were as follows: M1: 1.0 mm anterior to the bregma, 1.2 mm lateral to the brgma, and 0.8 mm ventral to the brain surface; S1: 2.0 mm lateral to the bregma, 0.5 mm ventral to the brain surface; and CPu: 0.5 mm anterior to the bregma, 2.0 mm lateral to the bregma, and 2.5 mm ventral to the brain surface. The mice were recovered from anesthesia with an intraperitoneal injection of atipamezole (1.5 mg/kg; Antisedan, Zenoaq) and maintained under regular health checks for one to six weeks.

Virus injection into marmoset brains was performed as described previously (*58*). All surgical procedures were conducted under aseptic conditions. Animals were anesthetized with intramuscular injections of ketamine (15 mg/kg) and medetomidine (50 μg/kg) and pre-medicated with intramuscular injections of atropine (40 μg/kg; Nipro ES Pharma), ampicillin (25 mg/kg; Viccillin, Meiji Seika pharma), and dexamethasone (80 μg/kg; Decadron, Aspen Japan), as well as a subcutaneous injection of a lactated Ringer’s solution (10 mL/kg; Solulact, Terumo) at 37°C. The animals were placed under deep anesthesia with isoflurane (1–2% in oxygen, Pfizer) inhalation. The head was fixed to a stereotaxic apparatus (SR-6C-HT, Narishige). Heart rate, pulse oxygen (SpO_2_), and rectal temperature were continuously monitored. A small hole was made in the skull with a dental drill. A glass micropipette with a tip diameter of 50 μm was filled with the viral solution (AAV2/1-SynTetOff EGFP-APEX2-BGHpA, 1.32 × 10^11^ IFU/mL). After incision of the dura, the pipette was slowly lowered to the target depth and fixed for 3 min. 0.15 μl of the viral solution was injected at a rate of 75 nl/min with a microsyringe pump (Legato 130, KD Scientific). The micropipette was held in place for 5 min and then extracted. The injection coordinates were as follows: 9.25 mm, 8.2 mm, 7.2 mm, and 6.15 mm anterior to the interaural line, 5.0 mm lateral to the midline, and 1.0 mm ventral to the brain surface. After the topical administration of gentamicin (Nichi-Iko Pharmaceutical), the head skin was closed by suturing. The animals were then received intramuscular injections of dexamethasone (80 μg/kg), diclofenac sodium (1.0 mg/kg, 11147700J1057, Novartic Japan), and ampicillin (25 mg/kg), as well as a subcutaneous injection of lactated Ringer’s solution (10 mL/kg) at 37°C. After surgery, the animals were recovered from anesthesia with an intramuscular injection of atipamezole (40 to 480 μg/kg), and ampicillin was administered for two days (25 mg/kg/day). The animals were maintained under regular health checks for six weeks.

#### DAB-Ni^2+^ labeling by APEX2

The effects of Sca*l*eSF clearing on its peroxidase activity of APEX2 were assessed by the polymerization of DAB. The brains were fixed with 4% PFA containing 0.2% GA seven to ten days after the injection of the AAV2/1-SynTetOff-EGFP-APEX2-BGHpA vector into the S1. The brain slices (1-mm thick) expressing EGFP-APEX2 fusion proteins were cleared with Sca*l*eSF. EGFP fluorescence in the slices was examined under the fluorescence stereomicroscope. After deSca*l*ing with PBS(–), the slices were cryoprotected in 30% sucrose in 0.1 M PB at 4°C, embedded in OCT compound (4583, Sakura Finetek), and frozen in liquid nitrogen-cooled isopentane. The slices were cut into 40-μm-thick sections on a freezing microtome (SM2000R, Leica Microsystems). Following CLSM imaging, the sections were permeabilized with PBS containing 0.3% Triton X-100 (0.3% PBS-X) and developed in 0.05% DAB.4HCl (347-00904, Dojindo), 25 mM nickel ammonium sulfate (24217-82, Nacalai Tesque), and 0.0003% H_2_O_2_ in 50 mM Tris-HCl (pH 7.6) (DAB-Ni^2+^ solution).

#### DAB-Ni^2+^ labeling by APEX2/BT-GO reaction

DAB polymerization in brain slices cleared with Sca*l*eSF was enhanced with APEX2/BT-GO (biotinylated tyramine-glucose oxidase) reaction, in which biotin molecules were deposited with tyramide signal amplification (TSA) reaction using its peroxidase activity of APEX2. Brains were fixed with 4% PFA containing 0.2% GA and cut into 1-mm-thick slices. The expression of EGFP-APEX2 in brain slices was examined as described above. The slices were then permeabilized for 4 hr with 0.2% PBS-X containing 2% bovine serum albumin (BSA) (01863-77, Nacalai Tesque), washed thrice with 0.1 M PB, and incubated for 4 hr in a BT-GO reaction mixture that contained 25 μM biotinylated tyramine and 3 μg/mL glucose oxidase (16831-14, Nacalai Tesque) (*42, 57, 59*). TSA reaction was initiated by adding 2 mg/mL of β-D-glucose (16804-32, Nacalai Tesque) into the reaction mixture and it proceeded for 2 hr. The brain slices were washed with PBS(–), fixed with 4% PFA in 0.1 M PB overnight at 4°C, and cleared with Sca*l*eSF. Cryosections (40- or 50-μm thick) or vibratome sections (50-μm thick) were prepared from deSca*l*ed slices as described above. Some of the sections were counterstained with DAPI (1 μg/ml, D1306, Thermo Fisher Scientific) in PBS for 2 hr on ice. The sections were then reacted with avidin-biotinylated peroxidase complex (ABC) (1:50 diluted in PBS, PK-6100, Vector Laboratories) in PBS containing 2% BSA overnight at 4°C and developed in DAB-Ni^2+^ solution on ice. CLSM imaging was performed prior to the ABC reaction.

#### Bright-field microscopy

Bright-field images of tissue sections were obtained with a light microscope (BX-51, Olympus) equipped with dry objectives (10× UPlanApo, NA = 0.40, WD = 3.1 mm, Olympus; 40× UPlanApo, NA = 0.85, WD = 0.2 mm, Olympus) and a CCD camera (DP72 or DP74, Olympus). DAB-Ni^2+^-labeled sections were mounted onto glass slides (Superfrost micro slide glass APS-coated, Matsunami Glass) and coverslipped with 50% glycerol, 2.5% 1,4-diazabicyclo[2.2.2]octane (DABCO) (049-25712, Wako Pure Chemical Industries), and 0.02% sodium azide (31233-55, Nacalai Tesque) in PBS.

#### Image processing

Three-dimensional renderings of CLSM image stacks were created with Imaris software (ver. 9.0.0, Bitplane). Images appearing in Fig. 1F–J were deconvoluted with Huygens Essential software (ver. 18.10.0p8, Scientific Volume Imaging) before the rendering process. Maximum intensity projection and orthogonal images were created using Leica Application Suite X (LAS X, ver. 3.5.5.19976, Leica Microsystems) and Imaris software. Three-dimensional reconstruction of FIB-SEM datasets was conducted using Dragonfly software (ver. 2020.2.0.941, Object Research System). The global brightness and contrast of the images were adjusted with ImageJ, Adobe Photoshop CS6 (Adobe), and CANVAS X DRAW.

#### Statistical analysis

Data are represented as means ± standard deviations (SDs). The exact values of n are indicated in the corresponding figure legends. For comparisons between groups, unpaired Student’s t-test was used (Fig. 1D). For comparisons among independent groups, one-way analysis of variance (ANOVA) (Fig. 2B), Kruskal–Wallis test followed by Steel–Dwass *post hoc* test (Fig. 3B), or Kruskal–Wallis test (Fig. S3C) was used. For comparisons between groups over time, two-way repeated measures ANOVA followed by Tukey *post hoc* test was used. The equality of probability distributions was assessed using Kolmogorov–Smirnov test. All tests were two-sided. Statistical analyses were conducted using EZR (ver. 1.41, Saitama Medical Center, Jichi Medical University) (*60*) and GraphPad Prism 8 (GraphPad Software). Statistical significance was set at P < 0.05.

## Acknowledgments

The authors thank Drs Atsushi Miyawaki and Hiroshi Hama (RIKEN Center for Brain Science) for their critical reading of the manuscript and constructive advice, and Editage (https://www.editage.com) for English language editing. We are also grateful to Keiko Okamoto-Furuta and Haruyasu Kohda (Division of Electron Microscopic Study, Center for Anatomical Studies, Graduate School of Medicine, Kyoto University) for technical assistance in electron microscopy.

## Author contributions

T.F., K.Y., and H.H. designed the study; T.F., K.Y., K.I., S.O., M.T., S.K., Y.I., and H.H. executed the experiments; T.F., K.Y., M.T., S.K., Y.I., A.T., and H.H. analyzed the data; T.F., K.Y., and H.H. wrote the original draft; T.F., K.Y., A.Y., Y.U., M.K., T.I., and H.H. edited the manuscript. All authors discussed the results and concurred on the contents of this manuscript.

## Data and materials availability

The datasets generated during and/or analyzed during the current study and all biological materials reported in this article are available from the corresponding author on reasonable request.

