## Supplementary Figures for "Title: Multi-Scale LM/EM Neuronal Imaging from Brain to Synapse with a Tissue Clearing Method, Sca*l*eSF"

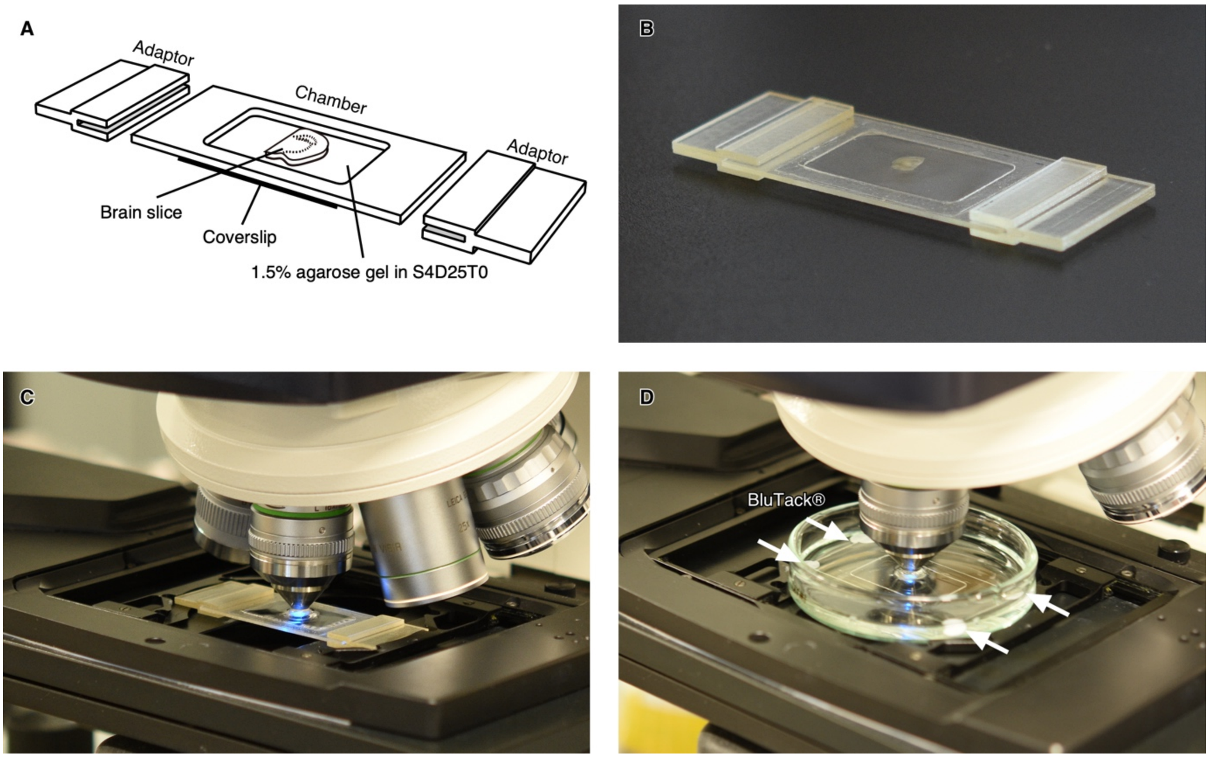


**Fig. S1. Customizable 3D-printed imaging chambers for Sca*l*eSF-treated tissues.** (**A**, **B**) A schematic drawing (A) and a picture (B) of a customizable 3D-printed imaging chamber. The imaging chamber comprises a chamber frame, a bottom coverslip, and microscope stage adaptors. Brain slices cleared with Sca*l*eSF are mounted on the bottom coverslip and embedded in 1.5% agarose gel in Sca*l*eS4D25(0) solution (Sca*l*eS4 gel). The frame and adaptors can be customized according to the size and thicknesses of brain slices. The bottom coverslip allows for imaging with inverted microscopes. (**C**, **D**) Imaging setups with the imaging chamber and an upright CLSM. (**C**) The imaging chamber mounted on a microscope stage using the microscope stage adaptors. (**D**) The imaging chamber without the adaptors is immersed in Sca*l*eS4 solution and mounted on the microscope stage. Blu-tack® is used to hold the imaging chamber firmly to the glass dish. Note that brain slices in (B–D) are not optically cleared.

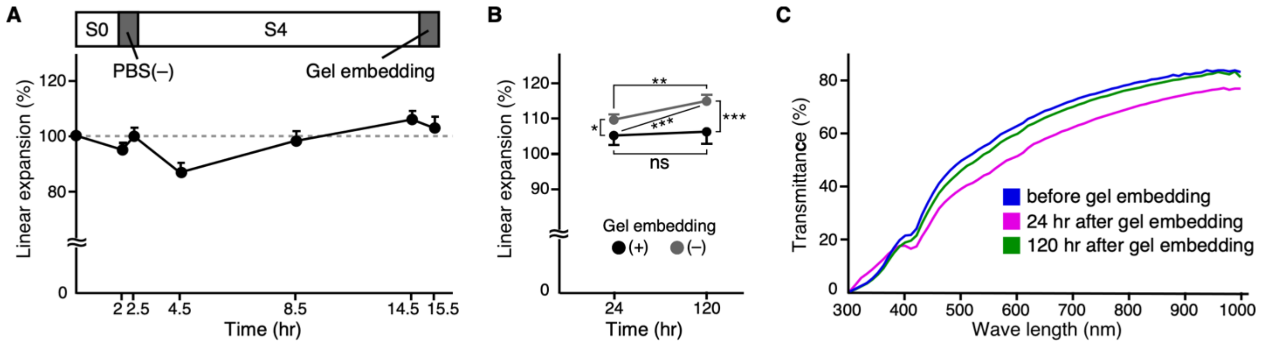


**Fig. S2. Changes in size and transparency of the mouse brain slices after embedding with Sca*l*eS4 gel.**

(**A**) Changes in the size of mouse brain slices during Sca*l*eSF treatment (n = 3 brain hemispheres). (**B**) Changes in the size of Sca*l*eSF-treated mouse brain slices with or without Sca*l*eS4 gel embedding (n = 3 brain hemispheres each; gel embedding, $F_{1,4} = 34.2$, $P = 0.0043$; time, $F_{1,4} = 83.45$, $P = 0.0008$; interaction, $F_{1,4} = 42.38$, $P = 0.0029$; two-way repeated measures ANOVA; * $P < 0.05$, ** $P < 0.01$, *** $P < 0.001$; Tukey *post-hoc* test). (**C**) Transmission curves of Sca*l*eSF-treated mouse brain slices before, and 24 hr and 120 hr after Sca*l*eS4 gel embedding (n = 3 brain hemispheres each). The thickness of brain slices is 1 mm. Error bars represent SDs.

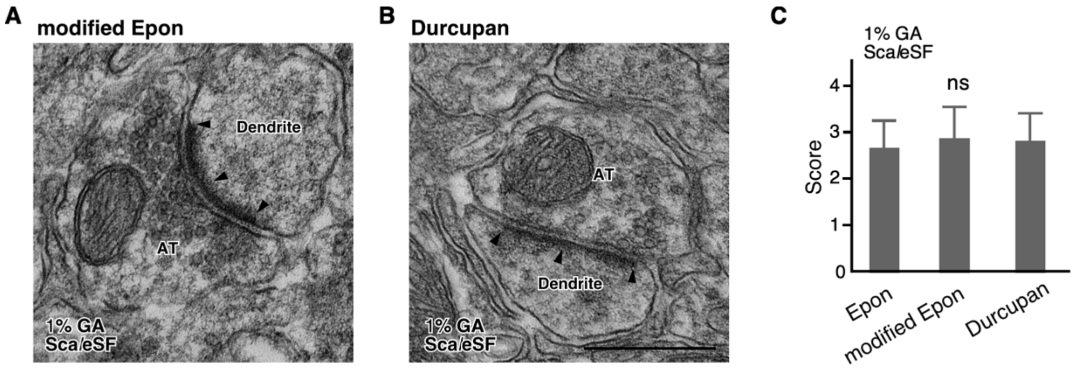


**Fig. S3. Effects of different epoxy resin embedding methods on ultrastructure preservation in Sca*l*eSF-treated mouse brain slices.**

(**A**, **B**) TEM images of Sca*l*eSF-treated mouse brain slices embedded with the modified Epon method (A) or Durcupan (B). When using the modified Epon method, resin-polymerization is initiated after pre-incubation in an epoxy mixture without an accelerator. The brains are fixed with 4% PFA containing 1% GA. (**C**) Scoring of membrane continuity of the presynaptic terminals for each condition. Over 90%, 50–90%, 10–50%, and less than 10% membrane continuity of the presynaptic terminals are scored as 4, 3, 2, and 1, respectively. There are no significant differences among the groups (n = 34 synapses, Epon; n = 31 synapses, modified Epon; n = 31 synapses, Durcupan; n = 3 mice for each condition; $H = 1.583$, $df = 2$, $P = 0.4532$, Kruskal–Wallis test). Error bars represent SDs. Scale bar: 500 nm.

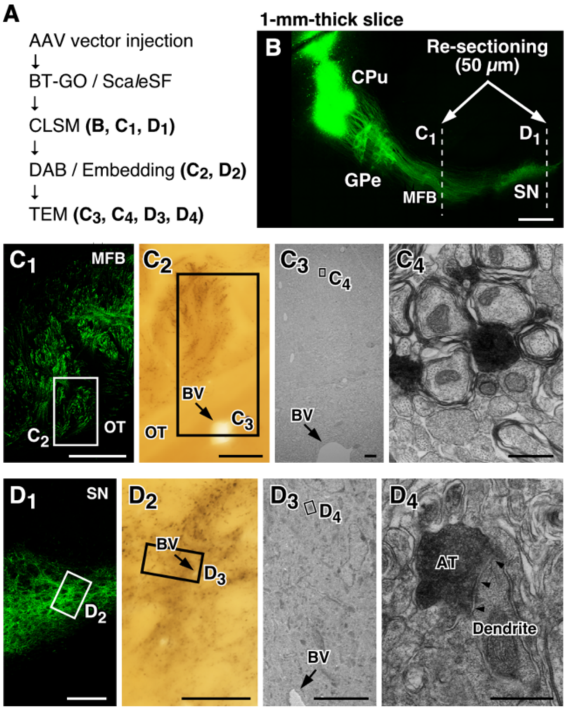


**Fig. S4. Multi-scale LM/EM neuronal imaging of the mouse striatonigral pathway.**

(**A**) The procedure of multi-scale LM/EM neuronal imaging of the mouse striatonigral pathway. (**B**) A maximum intensity projection image of the striatonigral pathway labeled with the AAV2/1-SynTetOff-EGFP-APEX2-BGHpA vector. A 1-mm-thick parasagittal brain slice is cleared with Sca*l*eSF four weeks after injection of the virus into the CPu. Sections of 50-μm thickness are cut along dotted lines. (**C**, **D**) Correlated fluorescent ($C_1$, $D_1$), bright-field ($C_2$, $D_2$), and TEM images ($C_3$, $C_4$, $D_3$, $D_4$) at the level of MFB (C) and SN (D). (**$C_1$**, **$D_1$**) CLSM imaging. (**$C_2$**, **$D_2$**) DAB-$Ni^{2+}$ labeling with APEX2/BT-GO reaction in the rectangle in ($C_1$) and ($D_1$). (**$C_3$**, **$D_3$**) A TEM image of the rectangle in ($C_2$) and ($D_2$). (**$C_4$**, **$D_4$**) A high magnification image of the rectangle in ($C_3$) and ($D_3$). Arrows in ($C_2$, $C_3$) and ($D_2$, $D_3$) indicate the identical blood vessel, respectively. Arrowheads in ($D_4$) point to a postsynaptic membrane. AT, axon terminal; BV, blood vessel. Scale bars: 500 μm in (B), 200 μm in ($C_1$, $D_1$), 50 μm in ($C_2$, $D_2$), 10 μm in ($C_3$, $D_3$), and 500 nm in ($C_4$, $D_4$).
